# The microbiome affects liver sphingolipids and plasma fatty acids in a murine model of the Western diet based on soybean oil

**DOI:** 10.1101/2020.09.03.281626

**Authors:** Sara C. Di Rienzi, Elizabeth L. Johnson, Jillian L. Waters, Elizabeth A. Kennedy, Juliet Jacobson, Peter Lawrence, Dong Hao Wang, Tilla S. Worgall, J. Thomas Brenna, Ruth E. Ley

## Abstract

Studies in mice using germfree animals as controls for microbial colonization have shown that the gut microbiome mediates diet-induced obesity. Such studies use diets rich in saturated fat, however, Western diets in the USA are enriched in soybean oil, composed of unsaturated fatty acids (FAs), either linoleic or oleic acid. Here we addressed whether the microbiome is a variable in fat metabolism in mice on a soybean oil diet. We used conventionally-raised, low-germ, and germfree mice fed for 10 weeks diets either high (HF) or low (LF) in high-linoleic-acid soybean oil as the sole source of fat. All mice, including germfree, gained relative fat weight and consumed more calories on the HF versus LF soybean oil diet. Plasma fatty acid levels were generally dependent on diet, with microbial colonization status affecting *iso*-C18:0, C20:3n-6, C14:0, and C15:0 levels. Colonization status, but not diet, impacted levels of liver sphingolipids including ceramides, sphingomyelins, and sphinganine. Our results confirm that absorbed fatty acids are mainly a reflection of the diet, and show that microbial colonization influences liver sphingolipid pools.

## 1. Introduction

The microbiome has emerged as central to the health of an organism[1]. Germfree animals, animals devoid of a microbiome, often serve as controls to a understand a variety of host phenotypes, from metabolism, the development of the immune system, the structure and function of gut cells, to animal behavior [2,3]. Early studies indicated that germfree mice are protected from diet induced obesity[4–7]. Subsequent studies observed that this protection is diet-dependent: germfree mice on a lard or tallow-based diet are protected, while those on a palm or coconut oil diet are not[8,9]. Although the specific fat used in these studies differ, they constitute predominantly saturated fat diets, which are not wholly representative of the modern Western diet. Soybean oil is the major oil consumed in the United States of America[10] and contains over 85% unsaturated omega-6 fatty acids (FAs). Unsaturated and saturated fats are observed to differentially impact the host’s weight, metabolism, fat deposition, and immune system[11–13]. The interaction of microbiome with a predominately unsaturated omega-6 fatty acid fat diet remains to be explored.

Here, we address the effect of a soybean oil diet on germfree, low-germ, and conventional mice with regards to weight, fat gain, circulating plasma lipids, and hepatic sphingolipids. We fed mice two paired soybean oil diets differing in calories from fat, with all of the fat deriving from high linoleic acid soybean oil (SBO), from weaning for 10 weeks. At the end of the 10-week period, we gavaged (oral delivery to the stomach) the animals with a large bolus of linoleic acid (C18:2n-6, LA), alpha-linolenic acid (C18:3n-3, ALA), or PBS control to observe potential microbial and host metabolic interactions with two FAs present in SBO. Following absorption of these FAs into the bloodstream, we assessed fat pad mass, circulating plasma FAs, and hepatic sphingolipids. Our results detect no effect of gut microbial colonization status on relative fat gain, which was higher for HF versus LF animals. Higher relative fat gain in HF fed animals correlated with increased food intake for HF versus LF fed animals in colonized mice, though not in GF mice. For both plasma FAs and hepatic sphingolipids, we observe that the presence of a microbiome as well as diet affected lipid pools. As expected for plasma FAs, diet was a greater contributor than microbial colonization status to the lipid pools. Conversely, for hepatic sphingolipid levels, microbial colonization status was a better predictor than diet. Our results indicate that microbes weakly affect lipids prior to absorption but have a greater effect on downstream lipid pathways in a host on a soybean oil diet.

## 2. Materials and Methods

### 2.1 Mouse experiments

All animal experimental procedures were reviewed and approved by the Institutional Animal Care and Usage Committee of Cornell University protocol 2010-0065. We used three sets of conventional male C57BL/6 mice (n=32, 35, 24) and two sets of germfree male C57BL/6 mice (n=36, 28) (**Table 1**). At weaning (3 weeks of age), littermates were split into cages with up to four mice/cage. Littermates were split to balance mouse weights within a cage and between the two diets. Conventional mice were maintained in the Accepted Pathogen Facility at Cornell University with filter top cages and the germfree mice in flexible, plastic (“bubble”) isolators[3] at Cornell University. All animals within a given germfree study were maintained within the same isolator at the same time. Animals in all studies were maintained under a 12-hour light cycle.

**Table 1.** Mouse groups.

| Study | Microbial status | Genotype | N | Vendor | N for plasma FAs | N for hepatic sphingolipids | N for qPCR |
| --- | --- | --- | --- | --- | --- | --- | --- |
| CNV1 | Conventional (MURINE Pathogen Free) | C57BL/6J | 32 | F2 generation* | 28 | 9 | 22 |
| CNV2 | Conventional (MURINE Pathogen Free) | C57BL/6NTac | 35 | Taconic Biosciences (Hudson, NY, USA) | 34 | 12 | 10 |
| CNV3 | Conventional (MURINE Pathogen Free) | C57BL/6NTac | 24 | Taconic Biosciences (Hudson, NY, USA) | 24 | 0 | 0 |
| GF (GF1) | Germfree | C57BL/6NTac | 36 | Taconic Biosciences (Hudson, NY, USA) | 35 | 12 | 28 |
| LG (GF2) | Low germ | C57BL/6NTac | 28 | Taconic Biosciences (Hudson, NY, USA) | 27 | 8 | 25 |

\* bred in the Accepted Pathogen Facility at Cornell University, originally from Jackson Laboratories (Bar Harbor, ME, USA)

The cages were supplied with either the LF (low fat, 16% kcal SBO) or HF (high fat, 44% kcal SBO) diet (**Table 2**) *ad libitum*. Diets were custom designed by Envigo (formerly Harlan Laboratories, Inc., Madison, WI, USA) and delivered pelleted, irradiated, and vacuum packed. For the conventional mice, we stored open, in-use diet bags at 4°C and unopened, bags at −20°C. For germfree mice, the in-use bags were stored within the germfree isolator where the animals were housed. We stocked cages with Pure-o-cel (The Andersons, Maumee, Ohio, USA), cotton nestlets, and plastic houses so to avoid the introduction of exogenous fat. For the germfree mice, all supplies were autoclaved at conditions to render the supplies germfree[3]. To bring the food into the germfree isolator, the vacuum-packed food bags were soaked in Clidox-S (Pharmacal Research Laboratories, CT, USA) or Exspor (Ecolab Inc., MN, USA) for least 30 minutes before moving the bags into the inner-port door where they were fumigated with Clidox-S or Exspor for a minimum of 2 hours. For all mice, food pellets were placed in the cages and not on the wire racks to minimize loss and crumb buildup of the diets as the HF SBO diet does not maintain pelleted form well and turns to powder easily. Twice weekly, we completely replaced cages and food. We weighed the amount of new food provided. To obtain mouse weights, we weighed mice in plastic beakers at the same approximate time of day twice weekly. To weigh germfree mice, we used a metal scale that was autoclaved and bought into the isolator.

**Table 2.**
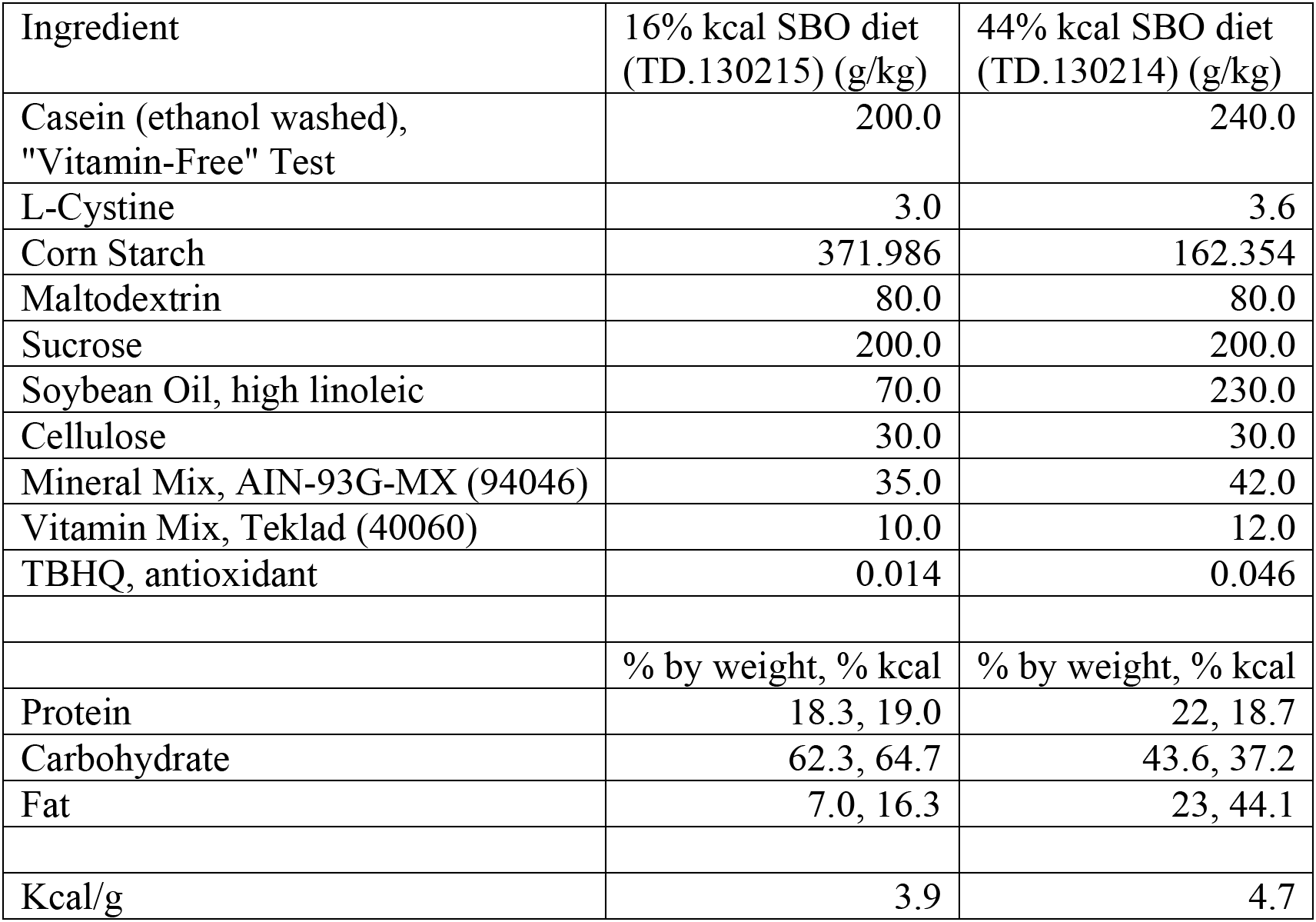
Soybean oil mouse diets.

| Ingredient | 16% kcal SBO diet<br>(TD.130215) (g/kg) | 44% kcal SBO diet<br>(TD.130214) (g/kg) |
| --- | --- | --- |
| Casein (ethanol washed),<br>"Vitamin-Free" Test | 200.0 | 240.0 |
| L-Cystine | 3.0 | 3.6 |
| Corn Starch | 371.986 | 162.354 |
| Maltodextrin | 80.0 | 80.0 |
| Sucrose | 200.0 | 200.0 |
| Soybean Oil, high linoleic | 70.0 | 230.0 |
| Cellulose | 30.0 | 30.0 |
| Mineral Mix, AIN-93G-MX (94046) | 35.0 | 42.0 |
| Vitamin Mix, Teklad (40060) | 10.0 | 12.0 |
| TBHQ, antioxidant | 0.014 | 0.046 |
|  | % by weight, % kcal | % by weight, % kcal |
| Protein | 18.3, 19.0 | 22, 18.7 |
| Carbohydrate | 62.3, 64.7 | 43.6, 37.2 |
| Fat | 7.0, 16.3 | 23, 44.1 |
| Kcal/g | 3.9 | 4.7 |

We collected fresh fecal samples once weekly from the beakers into tubes on dry ice, which were later stored at −80°C. For the germfree mice, fecal samples were collected per cage and for the conventional mice, per mouse. The conventional mice were handled exclusively inside of a biosafety cabinet and we changed personal protective equipment and wiped all surfaces with a sterilant between cages to prevent cross-contamination.

Food consumption was determined twice weekly in half of the cages for the CNV1, CNV2, GF1, and LG (GF1, see below) experiments. We filtered food crumbs out of the used bedding using a large hole colander followed by a fine mesh sieve, weighed the recovered food, and subtracted this amount from the known amount of food provided.

After 10 weeks on the SBO diets, mice were gavaged orally with 6 mg per gram mouse weight PBS, LA, or ALA. Within a cage, we gavaged roughly half of the mice with a FA and the other half with PBS, selecting which mouse received which gavage so to balance mouse weights between gavage groups. Following gavage, we moved mice to a fresh cage supplied with water but lacking food. After 1.5 hours, we euthanized mice by decapitation. Epididymal fat pads were removed and promptly weighed. We collected trunk blood resulting from decapitation into EDTA coated tubes collected and stored them on ice within 1 hour. Tubes were spun at 900 rcf at 4°C for 10 minutes and plasma was collected and stored at −80°C.

### 2.2 FA extraction and detection from plasma

We added 125 mg of heptadecanoic acid (C17:0, 99+% pure, Sigma Chemicals, St. Louis, MO, USA) to the plasma as an internal standard for absolute quantification of extracted FAs. The amount of plasma used in the extraction was 200 μl or the maximum of plasma acquired. We extracted lipids using the Bligh and Dyer method[14]. FAs were converted to FA methyl esters (FAMEs) and measured by gas chromatography. FA methyl esters (FAMEs) were prepared using 14% BF_3_ in methanol, dried under N_2_, dissolved in hexane with butylated hydroxytouelene, and stored at −20°C. FAMEs were measured in triplicate on a Hewlett-Packard 5890 series II gas chromatograph with a flame ionization detector (GC-FID) using H_2_ as the carrier. Peak areas were measured using PeakSimple software (SRI Instruments, Menlo Park, CA, USA). These peak areas were corrected using an equal weight mixture of known FAs measured multiple times throughout the GC run. See Su et al.[15] for further details. To account for the variable plasma volume used, each corrected peak area was normalized by the average plasma volume used across all samples (140 μl). We converted these peaks areas to quantitative amounts using the known amount of spiked C17:0 internal standard. All samples were run at random on the GC-FID.

To accurately determine structural identity of each peak in GC spectrum, select plasma samples were analyzed by covalent-adduct chemical ionization on a Saturn 2000 ion trap mass spectrometer (Varian, Inc., Walnut Creek, CA, USA)[16]. A few peaks could not be resolved to a single FA due to co-migration: C18:1n-9/7/? and several of the identified conjugated linoleic (CLA) and alpha-linolenic acids (ClnA). Arbitrary designators (e.g. ClnA.7, ClnA.2) were used to name these unresolved FAs.

### 2.3 Sphingolipid extraction from the liver tissue

Liver samples were defrosted on ice and homogenized in 1 mL of PBS in tubes containing 1 mm zirconium beads (OPS Diagnostics, Lebanon, NJ, USA) on a Mini Bead Beater homogenizer (BioSpec products, Bartlesville, OK, USA). Protein concentrations of the liver PBS homogenate were determined using the Lowry method (BioRad, Hercules, CA, USA) and 400 μg protein of liver homogenate was loaded into 96-deep well plates for lipid extractions. Lipids were extracted by adding 450 μL of 1:1 dichloromethane:methanol, 50 μL of 10% diethylamine in methanol, and 50 μL of the internal standard C12 ceramide (d18:1/12:0) (Avanti Polar Lipids, Alabaster, AL, USA) to samples with continuous shaking overnight (12 hours) on a plate shaker. The next day, 900 μL of 1:1 dichloromethane:methanol was added to each sample and gently mixed on a rotating shaker for an hour. Samples were then spun at 1500 x g for 15 minutes to pellet cellular debris. The supernatants were transferred to a new 96-deep well plate and stored at −20°C prior to analysis by high performance liquid chromatography-mass spectrometry (HPLC-MS).

Sphingolipid abundance was measured using targeted mass spectrometry on an Agilent 1200 HPLC liked to an Agilent 6430 triple quadrupole mass spectrometer according to previous methods[17].

### 2.4 Quantitative real-time PCR

To assess the microbial load in the mice (see **Table 1**), we performed quantitative real-time PCR (qPCR) using primers targeting the 16S rRNA gene[18]. We extracted DNA from fecal pellets taken from mice at weeks 9 to 11 on the SBO diets using the PowerSoil DNA isolation kit (Mo Bio Laboratories, Carlsbad, CA, USA) following the manufacturer’s protocol, and each sample was eluted with 50 μl Solution C6. We ran 10 μl reactions using the LightCycler 480 platform and the SYBR Green I Master kit (Roche Diagnostics Corporation, Indianapolis, IN, USA): 2 μl of DNA, each qPCR primer at 500 nM, and 5 μl of SYBR Green I Master mix. Cycling conditions were 5 minutes at 95°C followed by 45 cycles consisting of 10 seconds at 95°C, 20 seconds at 56°C, and 30 seconds at 72°C after which fluorescence from SYBR Green was read. Cycle threshold (C_t_) values were calculated using the absolute quantification/2^nd^ derivative max function available on the LightCycler 480 software. All reactions were run in triplicate and the mean C_t_ values were used in subsequent calculations. One conventional mouse sample was run in all qPCR runs to serve as an internal standard and 16S rRNA gene copy numbers from a given run are reported relative to this sample in that run. This standard was within one C_t_ across the three qPCR runs in this study.

### 2.5 Statistical analyses on FA and sphingolipid profiles

We utilized adonis (PERMANOVA)[19] to investigate how the full set of FAs in the conventional or germfree mouse sets were influenced by the technical and experimental variables. The technical variables included: normalized sum of FAs extracted (continuous variable), plasma volume used in extraction (continuous variable), mouse study (factor, see **Table 1**), FA extraction A date (Bligh-Dyer extraction part I, factor), FA extraction B date (Bligh-Dyer extraction part II, factor), GC run date (factor). The experimental variables were diet (factor), cage (factor), gavage (factor), and microbial status (conventional vs germfree, factor). We determined the differences among samples by computing a distance matrix using the Bray-Curtis dissimilarity metric[20] using the phyloseq package[21]. As we were unable to model the distance matrix with all technical and experimental variables, we first modeled the data using the full set of technical variables: *Bray-Curtis distance matrix ~plasma volume * normalized sum of FAs * study * FA extraction date A * FA extraction date B * GC run date*, where the multiplication sign indicates additive (e.g. *GC run date + study*) and interactive effects (e.g. *GC run date:study*) among the variables as implemented in R[22] using the vegan package[19] with 10,000 permutations. After running the full model, the non-significant terms were removed with p <0.05 being considered significant. Then the experimental variables were added: + *diet * gavage * microbial status * study* cage*. This model was then reduced so to keep the residuals less than 0.05. Principal Coordinate Analysis plots were made using the phyloseq package[21] with the ordplot function and a t-distribution. For analysis of hepatic sphingolipids, we utilized the model *Bray-Curtis distance matrix ~ diet * microbial status * study * cage*.

To determine the specific FAs altered by different experimental conditions (diet, gavage, and microbial status), we first reduced the list of FAs to only those present in two of the three conventional studies and one of the two germfree studies. Then we utilized a linear mixed model to determine which of the FA amounts were significantly affected by the experimental conditions as compared to a null model. The null model was *normalized FA mass ~ normalized sum of FAs + plasma volume + (1|cage) + (1|GC run date) + (1|FA extraction date A)*, where the terms *cage, GC run date, FA extraction date A*, and *study* were handled as random effects and all others as fixed effects. We excluded *FA extraction date B* as it is directly correlated with *date A*. For the experimental model, we added the terms *diet*gavage*microbial status*(1|study)*. These models were run in R[22] using the lme4 package[23] with REML = FALSE and the control optimizer set to “bobyqa”. We used a likelihood ratio test to compare the experimental and null models for each FA, and p-values were corrected with a Bonferroni correction; corrected p-values <0.05 were considered significant. For each FA found significant, we reduced the experimental terms (*diet*gavage*microbial status*(1|study*) so that the reduced model was not significantly different (p <0.05) from the full experimental model as determined by a likelihood ratio test. The lsmeans function[24] was used to compute the reduced model estimates on a model lacking the terms *(1|GC run date) + (1|FA extraction date A)* to avoid overfitting the data. Estimated differences between groups (e.g. HF VS LF diets) were calculated with a Tukey’s correction for multiple testing (p<0.01 was used to call significance). For hepatic sphingolipids, the null model was *sphingolipid data ~ (1|cage)* and the experimental model added the following terms: *diet*microbial status*(1|study)*. We corrected p-values using the Benjamini and Hochberg False Discovery Rate[25] instead of a Bonferroni correction with significance called at p<0.05 due to the lower power in these data compared to that for the plasma FAs. To investigate the composition of ceramide pools in the liver, we compared the abundance of each ceramide (Cer (d18:1/16:0), Cer (d18:1/18:0), Cer (d18:1/20:0), Cer (d18:1/22:0), Cer (d18:1/24:0), and Cer (d18:1/24:1)) relative to all ceramides in our sphingolipid model above. All statistical analyses were computed in R[22]. All t-tests are two-tailed, two-sample t-tests.

## 3. Results

### 3.1 Germfree mouse studies contained negligible to low levels of microbes

Germfree mice maintained for prolonged periods of time on diets that cannot be autoclaved will inevitably acquire a microbiota. Visually, GF1 and GF2 mice appeared germfree, as evidenced by their grossly enlarged cecal size and over-abundance of bile following oil gavage[2]. Although the GF1 experiment mouse feces had 16S rRNA copies indistinguishable to that of the extraction and qPCR blanks (t-test, p>0.05), values measured for the GF2 experiment were greater (t-test, p<10^−4^) (**Fig 1**). Assuming the blanks had no bacteria, GF2 fecal pellets are estimated are to have 10^4^ - 10^5^ bacterial cells per extracted fecal pellet, three orders of magnitude lower than the conventional animals (t-test, p<10^−7^), which have an estimated ~10^8^ bacterial cells. Hence, while the GF1 animals can be considered germfree (hereafter, GF), the GF2 animals are “low-germ” and hereafter labeled LG.

**Figure 1.**
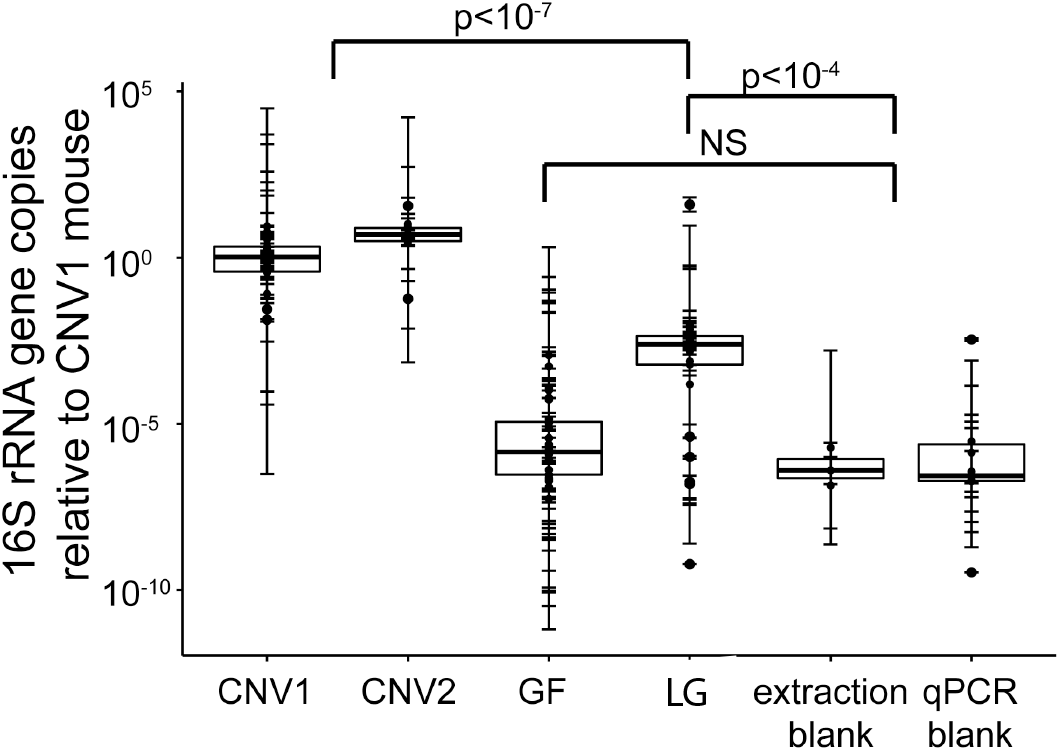
Microbial load in conventional and germfree mouse experiments. 16S rRNA gene copy number was determined by qPCR relative to a single mouse from the CNV1 experiment. For statistical analyses (t-test), the blanks were grouped together as were the conventional animals. Error bars show the standard deviations for each triplicate qPCR run, and box plots show the data distribution for each mouse study or blank group. N= 3 to 28 per group.

### 3.2 Fat mass gain is a function of SBO diet oil content, not microbial colonization status

To determine if GF and LG mice were able to accumulate more body fat on the HF diet than on the LF diet, we measured their weights over the course of the experiment (**Fig 2A** and **B**) and the mass of their epididymal fat pads at euthanasia (**Fig 2C**). Mice on the HF diet obtained greater epididymal fat pad mass compared to mice on the LF diet in each study, conventional, germfree, or low-germ (t-test, p-values <0.05, **Fig 2C, Table 3**). When comparing between the germfree and conventional studies, conventional animals on either diet had greater relative fat mass and greater epididymal fat pad mass compared to germfree or low-germ mice on the same diet (t-test, pvalues<10^−8^, **Fig 2B** and **C**). We measured food consumption in two of the three conventional studies and both of germfree studies (**Fig 2D**). From these data, we observed that mice on the HF diet consumed significantly more food, and more calories, compared to mice on the LF diet, except in the GF group, in which animals were only significantly different for consumed calories (t-test, p-values <0.05, **Table 3**). The GF germfree animals on the HF diet gained relatively more fat mass than those on the LF diet without increased food consumption likely due to the fact that the HF diet is more energy dense that the LF diet (**Table 2**).

**Figure 2.**
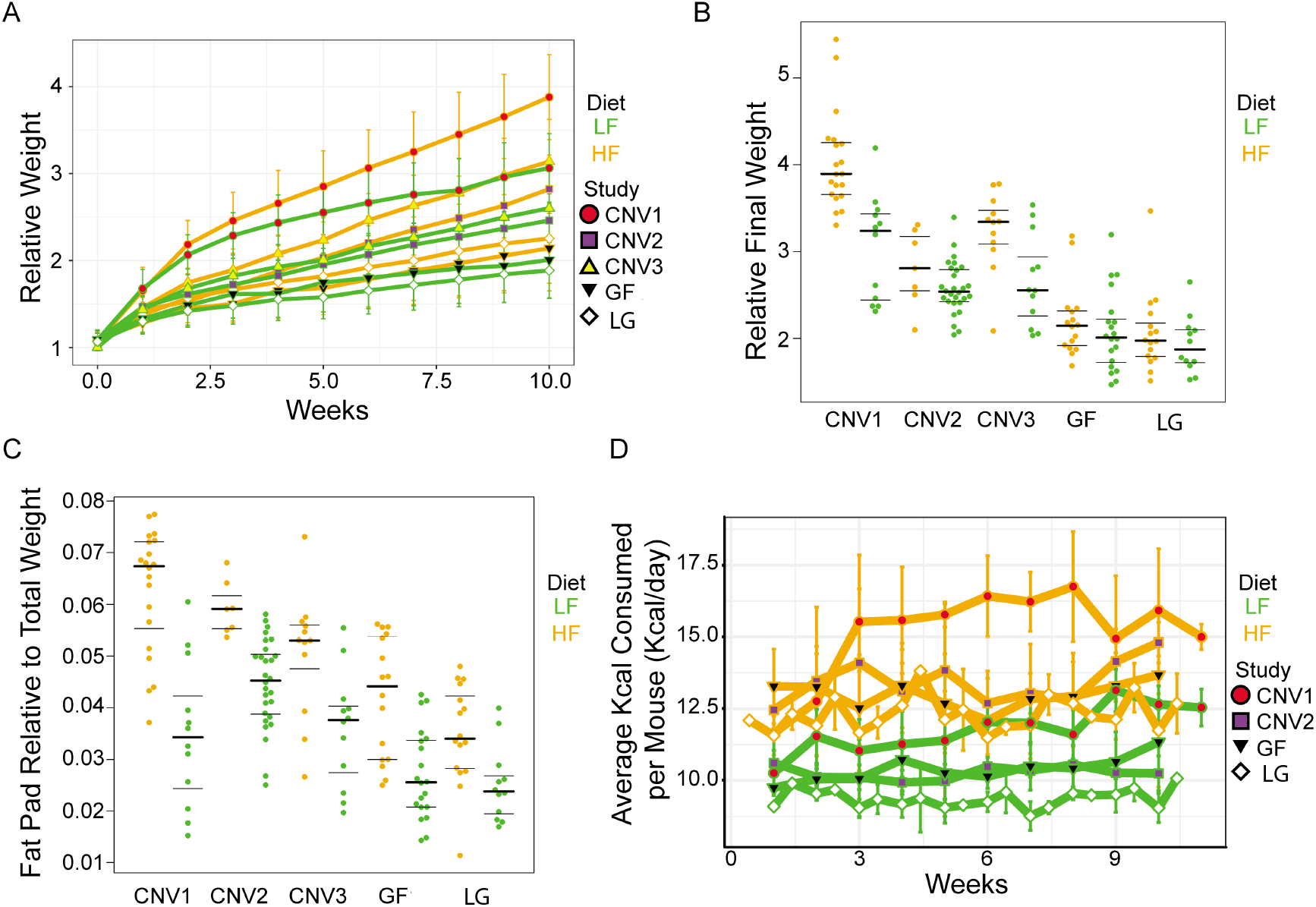
Mouse weight, fat pad mass, and food consumption on soybean oil diets. **A**. Weight of animals over time relative to weight at weaning when placed on SBO diets. **B**. Weight of animals at euthanasia relative to weight at weaning. **C**. Epididymal fat mass relative to total body weight at euthanasia. **D**. Average calorie consumption for each animal. Food consumption was measured per cage. For **A** and **D**, error bars indicate standard deviations across all animals in a study. For **B** and **C**, dark lines indicate the 50% quartile, and the two thinner lines show the 25% and 75% quartiles. See Table 3 for statistical analyses. Green = LF SBO diet; Orange = HF SBO diet; Red circles = CNV1; Purple squares = CNV2; Yellow upwards triangle = CNV3; GF = Black downwards triangle; White diamond = LG.

**Table 3.** Mouse weights and food consumption for each study.

|  |  |  |  |
| --- | --- | --- | --- |
| CNV1 |  |  |  |
| Diet | HF (n=20) | LF (n=12) | t-test p |
| Absolute Weight (g) | 36.8 | 31.8 | 0.0002645 |
| Relative Weight to Weaning Weight | 4.0 | 3.0 | 0.0001538 |
| Epididymal Fat Pad Absolute (g) | 2.3 | 1.1 | 5.992e-06 |
| Epididymal Fat Pad Relative to Total Weight | 0.063 | 0.035 | 1.298e-05 |
| Average Daily Food Consumption (g/day) | 3.22 | 3.02 | 0.01016 |
| Average Daily Kcal Consumption (Kcal/day) | 15.15 | 11.77 | 1.273e-13 |
| CNV2 |  |  |  |
| Diet | HF (7) | LF (28) | t-test p |
| Absolute Weight (g) | 35.7 | 34.5 | 0.07635 |
| Relative Weight to Weaning Weight | 2.8 | 2.6 | 0.2318 |
| Epididymal Fat Pad Absolute (g) | 2.2 | 1.6 | 0.0006235 |
| Epididymal Fat Pad Relative to Total Weight | 0.059 | 0.045 | 4.08e-05 |
| Average Daily Food Consumption (g/day) | 2.84 | 2.63 | 0.002529 |
| Average Daily Kcal Consumption (Kcal/day) | 13.37 | 10.27 | 4e-13 |
| CNV3 |  |  |  |
| Diet | HF (12) | LF (12) | t-test p |
| Absolute Weight (g) | 38.9 | 32.3 | 0.00302 |
| Relative Weight to Weaning Weight | 3.2 | 2.7 | 0.009278 |
| Epididymal Fat Pad Absolute (g) | 2.0 | 1.2 | 0.001867 |
| Epididymal Fat Pad Relative to Total Weight | 0.050 | 0.036 | 0.00542 |
| Average Daily Food Consumption (g/day) | ND | ND | ND |
| Average Daily Kcal Consumption (Kcal/day) | ND | ND | ND |
| GF (GF1) |  |  |  |
| Diet | HF (16) | LF (20) | t-test p |
| Absolute Weight (g) | 33.5 | 32.3 | 0.2209 |
| Relative Weight to Weaning Weight | 2.2 | 2.1 | 0.4115 |
| Epididymal Fat Pad Absolute (g) | 1.4 | 0.9 | 0.0007992 |
| Epididymal Fat Pad Relative to Total Weight | 0.042 | 0.027 | 0.0002224 |
| Average Daily Food Consumption (g/day) | 2.76 | 2.67 | 0.09075 |
| Average Daily Kcal Consumption (Kcal/day) | 12.99 | 10.40 | 1.33e-15 |
| LG (GF2) |  |  |  |
| Diet | HF (16) | LF (12) | t-test p |
| Absolute Weight (g) | 31.9 | 30.1 | 0.1634 |
| Relative Weight to Weaning Weight | 2.1 | 1.9 | 0.3592 |
| Epididymal Fat Pad Absolute (g) | 1.1 | 0.8 | 0.01076 |
| Epididymal Fat Pad Relative to Total Weight | 0.035 | 0.025 | 0.00511 |
| Average Daily Food Consumption (g/day) | 2.61 | 2.40 | 4.808e-08 |
| Average Daily Kcal Consumption (Kcal/day) | 12.28 | 9.36 | < 2.2e-16 |

### 3.3 Microbial status is a minor effector on plasma FAs

To investigate if the circulating FA profile differed according to microbial colonization status, we measured plasma FAs in all mice post-gavage with PBS, LA and ALA. We analyzed the entire set of measured FAMEs to determine which of the variables in our experimental design and sample handling influenced the data by using a Permutational Multivariate Analysis of Variance Using Distance Matrices (PERMANOVA) with the adonis function[19]. To do so, we created a distance matrix of the data using the Bray-Curtis dissimilarity metric[20], which computes the dissimilarity among individuals by composition and quantity. After model simplification, we were able to explain 97.2% of the data with the model shown in **Table 4**. These models indicate that diet (~13% of the data), microbial status (~5%), and gavage (~3%) contribute to plasma FAs.

**Table 4:** PERMANOVA on plasma FAs, residuals = 0. 02842.

| Term | R<sup>2</sup> (% variation) | p-value |
| --- | --- | --- |
| Normalized sum of FAs | 0.42505 | 0.0001 |
| Gavage:Cage | 0.16746 | 0.6625 |
| Diet | 0.12976 | 0.0001 |
| CNV, LG, or GF | 0.05638 | 0.0001 |
| Gavage | 0.03417 | 0.0001 |
| FA extraction B date | 0.02908 | 0.0001 |
| Plasma volume used in extraction:Study | 0.02879 | 0.0006 |
| Study:GC run | 0.02678 | 0.0016 |
| FA extraction A date | 0.02597 | 0.0001 |
| Study | 0.02467 | 0.0002 |
| Plasma volume used in extraction | 0.0139 | 0.0008 |
| Normalized sum of FAs:Plasma volume used in extraction | 0.00958 | 0.0035 |

Principal Coordinate Analysis (PCoA) plots also indicate that diet contributes to the set of plasma FAs, and microbial status less so (**Fig 3A** and **B**). Furthermore, a linear regression of total extracted plasma FAs and microbial load measured by qPCR showed no relationship (**Fig 4A**). These observations indicate that microbial status has a minor effect on the overall plasma FA profile, whereas the animal’s diet has a greater effect.

**Figure 3.**
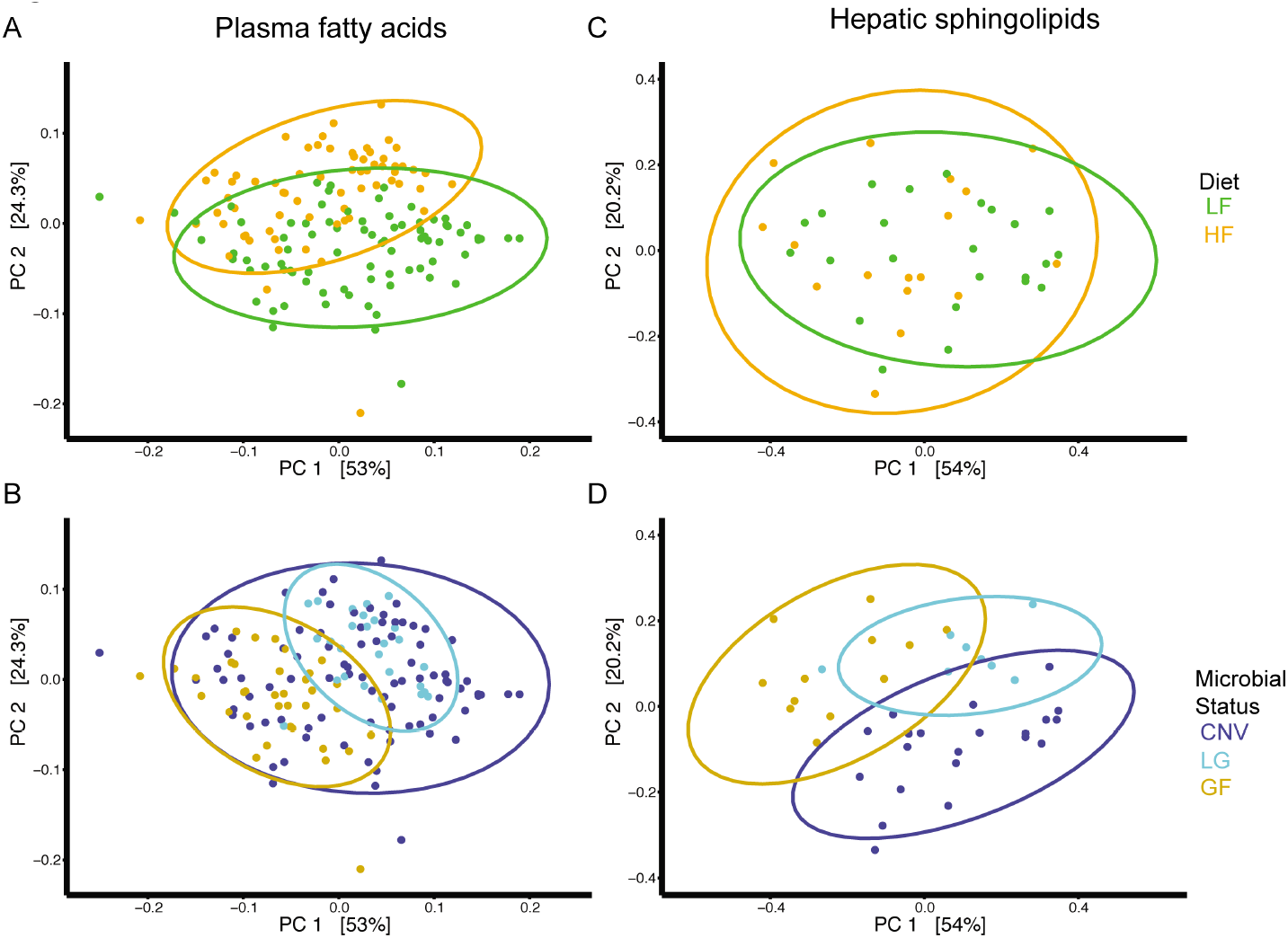
Total plasma FA compositions cluster by diet and microbial status whereas hepatic sphingolipid compositions cluster primarily by microbial status. PCoA plots from Bray-Curtis computed distance matrices for plasma FAs (**A,B**) and for hepatic sphingolipids (**C,D**), colored by diet (**A,C**) or microbial status (**B,D**). Green = LF SBO diet; Orange = HF SBO diet; Blue = CNV; Light blue = LG; Tan = GF.

**Figure 4.**
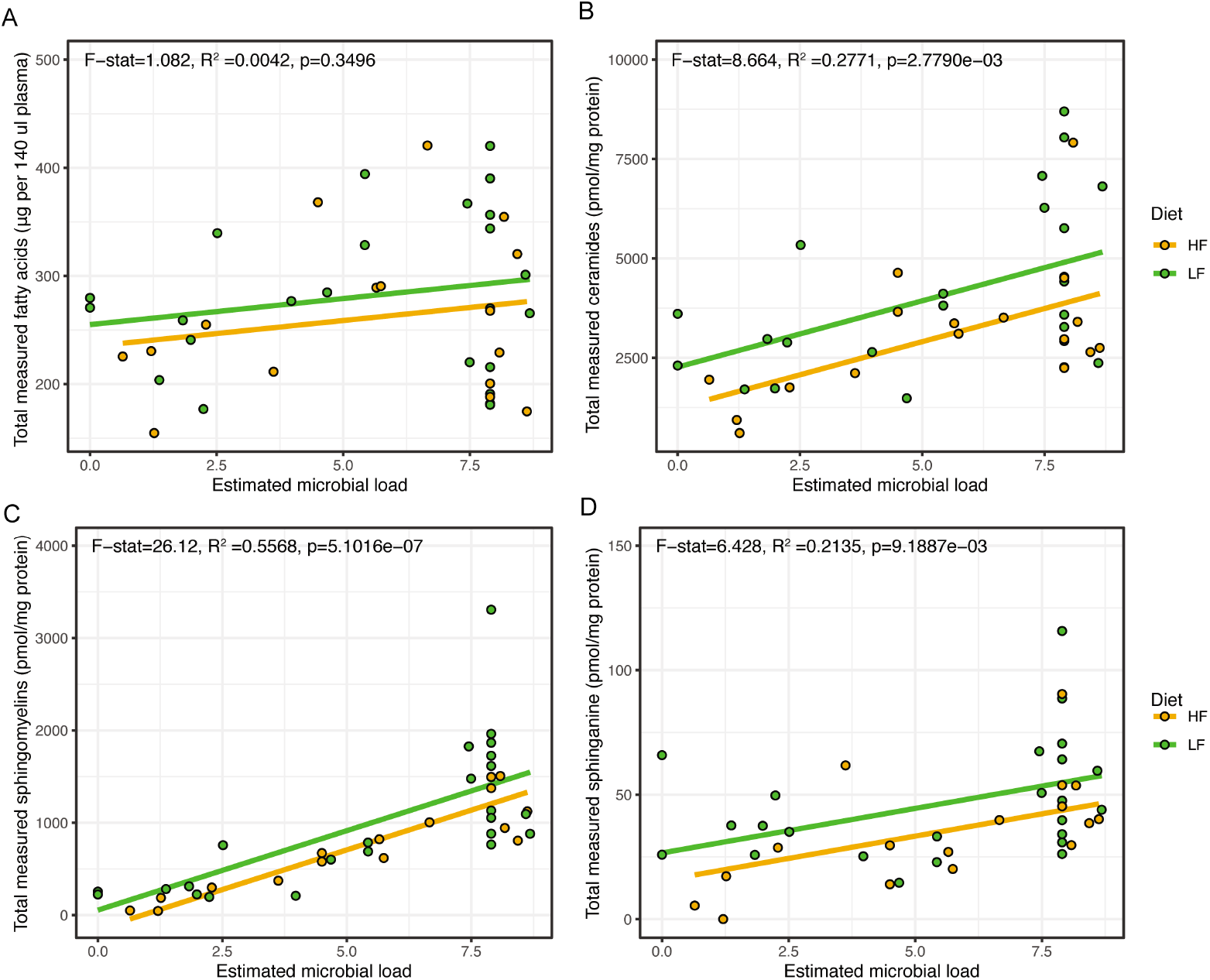
Hepatic sphingolipids not plasma FAs are correlated with microbial load. Estimated microbial load is derived from the average 16S rRNA copy number for all CNV mice, just GF1 mice (GF mice), or just GF2 mice (LG mice) qPCR on the 16S rRNA gene as reported in Figure 1. The data **A**, plasma FAs, **B**, hepatic ceramides, **C**, hepatic sphingomyelins, or **D**, hepatic sphinganine were fit using a linear regression. Only data from animals gavaged with PBS are shown to facilitate comparison between plasma FAs and hepatic sphingolipids. For the hepatic sphingolipids (**B**, **C**, and **D**), adjusted pvalues are shown, which were corrected using the Benjamini and Hochberg False Discovery Rate across all classes of sphingolipids. Only sphingolipid classes with adjusted p<0.05 are shown. Green = LF SBO diet; Orange = HF SBO diet.

### 3.4 Diet, gavage, and microbial status determine the quantity of specific plasma FAs

To determine which circulating FAs were impacted by diet, gavage, and microbial status we utilized a linear mixed model. After removing FAs not present in two of the three conventional studies and one of the two of the germfree studies (LG and GF mice), we considered 38 distinct FA peaks. Five of these 38 represent more than one FA that could not be adequately separated.

We identified 17 FAs with p <0.05 after a Bonferroni correction significantly impacted by diet and/or gavage (**Table 5**). Nine of the FAs were affected by diet, with the HF diet reducing the amount of the FA except for LA and ALA (**Tables 5**, **Supplementary Table 1**, and **Supplementary Figures S1-S5**), which were present in greater amounts in the high fat diet. As expected, the LA gavage increased plasma LA amounts (p-values<0.0001) and the amount of ALA was higher in animals gavaged with ALA (p-values<0.0001) (**Tables 5**, **Supplementary Table 1**, and **Supplementary Figures S3, S4**). Several ClnAs were also were higher when the animal had been gavaged with LA (p-values<0.001, **Tables 5, Supplementary Table 1**, and **Supplementary Figures S5, S6**). The interaction of diet and gavage was important for three FAs. In particular, only for the LF diet was C20:5n-3 present in higher amounts in animals gavaged with PBS (p-values<0.0001), and there was less C16:0 *iso* in HF-ALA mice compared with LF-ALA mice (p-values<0.01) (**Tables 5, Supplementary Table 1**, and **Supplementary Figures S1, S6**).

**Table 5.** Plasma FAs significantly impacted by diet, gavage, and/or microbial status.

| FA | Diet | Gavage | MS | Diet:Gavage | Diet:MS | Gavage:MS | Diet:Gavage:MS |
| --- | --- | --- | --- | --- | --- | --- | --- |
| C14:0 | LF><sup>1</sup>HF*** | PBS>ALA* | NS<sup>2</sup> |  | (HF)<sup>3</sup>GF>CNV*;<br>(HF)GF>LG*;<br>(CNV)LF>HF* |  |  |
| C15:0 |  | LA>ALA*;<br>LA>PBS* |  | (HF)LA>PBS* |  |  | (HF,LA)GF>CNV**;<br>(HF,GF)LA>ALA** |
| C16:0 *iso* | NS |  |  | (ALA)LF>HF* |  |  |  |
| C16:0 | LF>HF*** |  |  |  |  |  |  |
| C16:1n-7 | LF>HF*** |  |  |  |  |  |  |
| C18:0 *iso* | LF>HF*** |  |  |  | (CNV)LF>HF*** |  |  |
| C18:1n-7/9/? | LF>HF*** |  | NS |  |  |  |  |
| C18:2n-6 | HF>LF** | LA>ALA***;<br>LA>PBS*** | LG>CNV** |  |  |  |  |
| C18:3n-6 |  | PBS>LA<sup>5</sup>* | LG>CNV***;<br>LG>GF*** |  |  |  |  |
| C18:3n-3 | HF>LF* | ALA>LA***;<br>ALA>PBS*** |  |  |  | (CNV)ALA>LA***;<br>(CNV)ALA>PBS***;<br>(LG)ALA>LA***;<br>(LG)ALA>PBS***;<br>(GF)ALA>LA* |  |
| C20:3n-6 |  | PBS>LA<sup>5</sup>*** | CNV>GF** |  |  |  |  |
| ClnA.7 |  | NS | GF>CNV**;<br>GF>LG*** |  | (HF)GF>CNV**;<br>(HF)GF>LG*** | (LA)GF>CNV*;<br>(LA)GF>LG*** |  |
| C20:5n-3 |  |  |  | (LF)PBS>LA*** |  |  |  |
| ClnA.2 |  | LA>ALA***;<br>LA>PBS*** | GF>LG** |  |  |  |  |
| ClnAs<sup>4</sup> |  | LA>ALA**;<br>LA>PBS*** |  |  |  | (LA)GF>CNV*;<br>(LA)GF>LG***;<br>(GF)LA>ALA*;<br>(GF)LA>PBS*** |  |
| C22:5n-6 | LF>HF*** |  |  |  |  | NS |  |
| C24:1n-9 |  | ALA>LA<sup>5</sup>***;<br>PBS>LA<sup>5</sup>*** |  |  |  |  |  |

<sup>1</sup>FA is higher for condition on listed at left of “>”
<sup>2</sup>Term is required for the linear mixed model but term is not significant after multiple testing correction among contrasts.
<sup>3</sup>Effect listed is only for the indicated diet. For example, (HF)GF>CNV means that only for the HF diet was more FA present in GF compared to CNV animals.
<sup>4</sup>This peak represents two ClnAs.
<sup>5</sup>Model appears incorrect for estimate, see **Supplementary Figures**
\*, p<0.01; \*\*, p<0.001; \*\*\*, p<0.0001

Microbial status affected the amount of 10 FAs (**Tables 5**, **Supplementary Table 1**, and **Supplementary Figures S1-S6**). C20:3n-6 was the only FA only affected by microbial status and it was higher in conventional mice (p-values<0.001, **Figure 5, Tables 5**, **Supplementary Table 1**), suggesting it is produced by or its production in the host is stimulated by microbes. Similarly, C18:0 *iso* was higher in colonized mice, but only those on the LF diet (p-values<0.001, **Figure 5, Tables 5**, **Supplementary Table 1**). Other FAs were higher in GF mice: two ClnAs (ClnA.7 and ClnAs4) in GF animals gavaged with LA compared to CNV or LG mice, (p-values<0.01, **Tables 5**, **Supplementary Table 1, Supplementary Figures S5, S6**) and C14:0 and C15:0 in GF-HF diet animals (HF-LA animals for C15:0) (p-values<0.01, **Figure 5, Tables 5**, **Supplementary Table 1**), suggesting microbes directly or indirectly promote the metabolism of these FAs.

**Figure 5.**
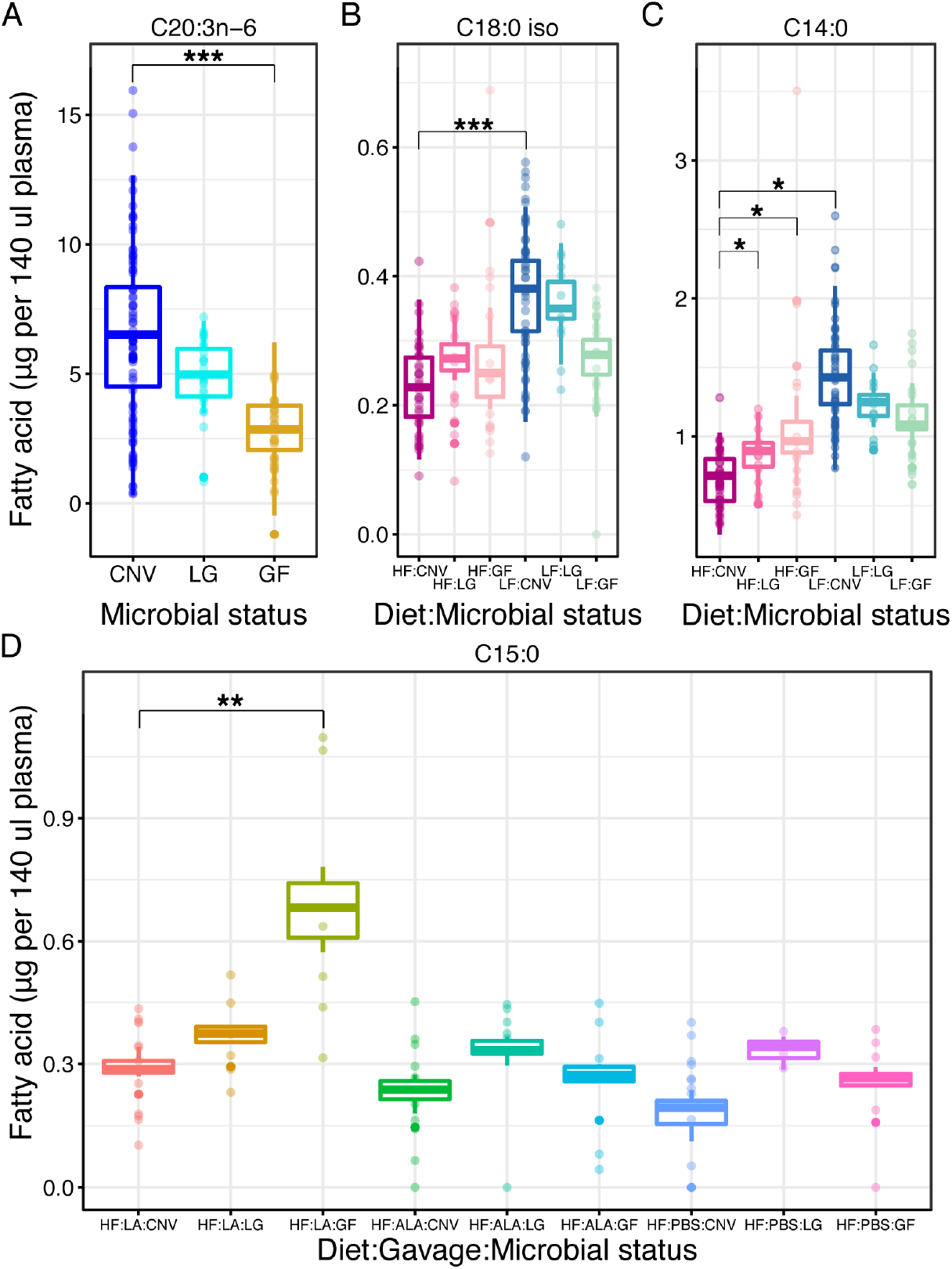
Plasma FAs differing by microbial load and diet. Plasma **A**, C20:3n-6, **B**, C18:0 *iso*, **C**, C14:0, and **D**, C15:0. Normalized data are shown as points and the boxplots show the covariate adjusted means from the least squares means estimates. N=3 to 148 per group. Blue = CNV; Light blue = LG; Tan = GF. Dark magenta = HF:CNV; Rose = HF:LG; Light pink = HF:GF; Navy = LF:CNV; Teal = LF:LG; Sea green = LF:GF; Red = HF:LA:CNV; Ochre = HF:LA:LG; Olive = HF:LA:GF; Green = HF:ALA:CNV; Jade = HF:ALA:LG; Maya blue = HF:ALA:LG; Cornflower purple = HF:PBS:CNV; Orchid = HF:PBS:LG; Lilac = HF:PBS:LG; Cherry pink = HF:PBS:GF.

### 3.5 Liver sphingolipids differ by microbial status

Hepatic lipids have been previously observed to be dependent on microbial colonization[26] and sphingolipids are known to be closely linked to metabolic diseases[27]. To determine if hepatic sphingolipids were affected by microbial state, we measured sphingolipids in livers of mice gavaged with PBS (see **Table 1**). Unlike observed for plasma FAs (**Fig 4A**), we observed that the total amounts of ceramides, sphingomyelins, and sphinganine were positively linearly related to the microbial load (**Fig 4B**). In accord, whereas the plasma FA data clustered readily by diet in PCoA (**Fig 3A**), the sphingolipid data did not (**Fig 3C**), and instead clustered well by microbial status (**Fig 3D**). The PERMANOVA analysis supported this observation: microbial status explained more of the variance in the sphingolipid data (~37%) compared to diet (~3%) (**Table 6**).

**Table 6.** PERMANOVA on hepatic sphingolipids, residuals = 0.01946.

| Term | R<sup>2</sup> (% variation) | p-value |
| --- | --- | --- |
| Cage | 0.53772 | 0.0001 |
| CNV, LG, or GF | 0.37423 | 0.0001 |
| Study | 0.03795 | 0.0003 |
| Diet | 0.03065 | 0.0008 |

To determine which specific sphingolipids were affected by diet and microbial status, we ran a linear mixed model. We observed six sphingolipids to vary by microbial status whereas diet contributed to only one sphingolipid in interaction with microbial status (**Table 7, Supplementary Table 2**, and **Figure 6**). Cer (d18:1/20:0), Cer (d18:1/24:1), Sa (d18:1), and SM (d18:1/24:1) mirror the trend observed for total sphingolipid amounts, whereby sphingolipids levels increase as microbial load increases (p-values<0.05, **Table 8, Supplementary Table 2**, and **Figure 6A-D**). Cer (d18:0/16:0) was lower in LG animals and Cer (d18:0/16:0) DH was higher in GF animals, but only those on the LF diet (p-values≤0.0001, **Table 8, Supplementary Table 2**, and **Figure 6E** and **F**).

**Table 7.** Hepatic sphingolipids significantly impacted by diet and/or microbial status.

| Sphingolipid | Diet | Microbial status | Diet:Microbial status |
| --- | --- | --- | --- |
| Cer (d18:1/16:0) |  | CNV>LG**; GF>LG*** |  |
| DHCer (d18:0/16:0) |  |  | (LF)GF>CNV***; (LF)GF>LG* |
| Cer (d18:1/20:0) |  | CNV>LG**; CNV>GF*** |  |
| Cer (d18:1/24:1) | NS<sup>1</sup> | CNV>LG*; CNV>GF*** |  |
| Sa (d18:0) |  | CNV>LG*; CNV>GF* |  |
| SM (d18:1/24:1) |  | CNV>LG**; CNV>GF*** |  |

Table presented as for **Table 5**.
\*, p<0.05; \*\*, p<0.001; \*\*\*, p<0.0001

**Figure 6.**
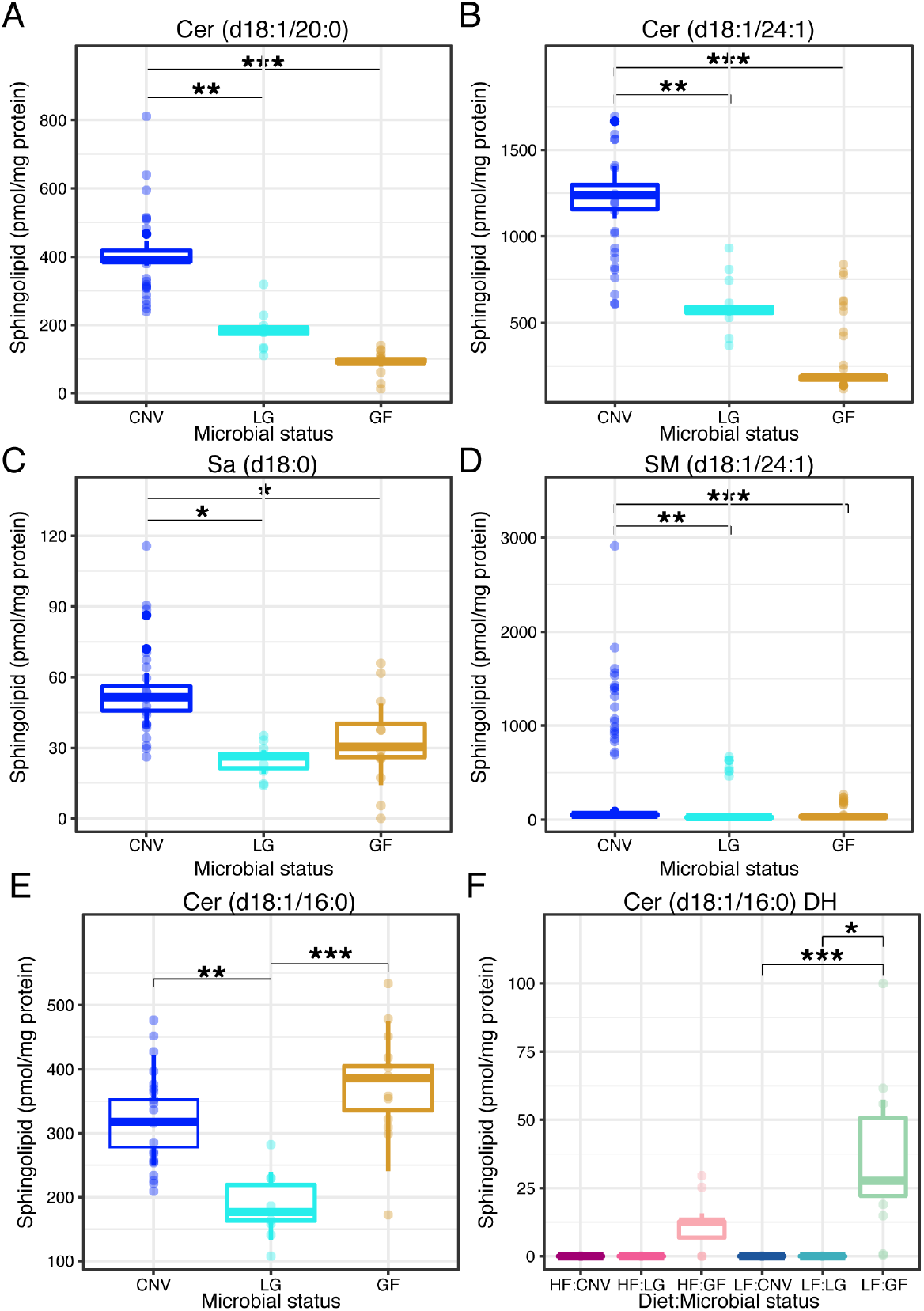
Hepatic sphingolipids differing by microbial load and diet. Hepatic **A**, Cer (d18:1/20:0), **B**, Cer (d18:1/24:1), **C**, Sa (d18:0), **D**, SM (d18:1/24:1), **E**, Cer (d18:1/16:0), and **F**, Cer (d18:1/16:0) DH. Data are plotted as in Figure 5. N=4 to 21 per group. Blue = CNV; Light blue = LG; Tan = GF. Dark magenta = HF: CNV; Rose = HF:LG; Light pink = HF:GF; Navy = LF:CNV; Teal = LF:LG; Sea green = LF:GF.

**Table 8.** Hepatic ceramide abundances significantly impacted by diet and/or microbial status.

| Ceramide | MS |
| --- | --- |
| Cer (d18:1/16:0) | GF>CNV***, GF>LG*** |
| Cer (d18:1/18:0) | GF>CNV*, GF>LG** |
| Cer (d18:1/20:0) | CNV>GF***, CNV>LG* |

Table presented as for **Table 5**.
\*, p<0.05; \*\*, p<0.001; \*\*\*, p<0.0001

Ceramide fatty acyl chain length is an important indicator of metabolic function. The Cer (d18:1/16:0)/Cer (d18:1/24:0) ratio is an indicator of metabolic syndrome with higher Cer (d18:1/16:0) levels producing diet induced obesity and insulin resistance[28]. In our data, we observed no significant differences in this ratio across all mice (p>0.05, data not shown). To better understand the composition of ceramide pools, we computed the abundance of each ceramide relative to the sum of all measured ceramides and ran the same linear mixed model used on all sphingolipids on this reduced set. Cer (d18:1/16:0) and Cer (d18:1/18:0) relative abundances were highest in the germfree animals (p-values<0.05, **Table 8, Supplementary Table 3**, and **Figure 7 A and B**) and Cer (18:1/20:0) was highest in the conventional animals (p-values<0.05, **Table 8, Supplementary Table 3**, and **Figure 7C**). These data illustrate that in soybean oil fed mice, hepatic ceramide abundance is altered by microbes, irrespective of the percentage of calories from soybean oil.

**Figure 7.**
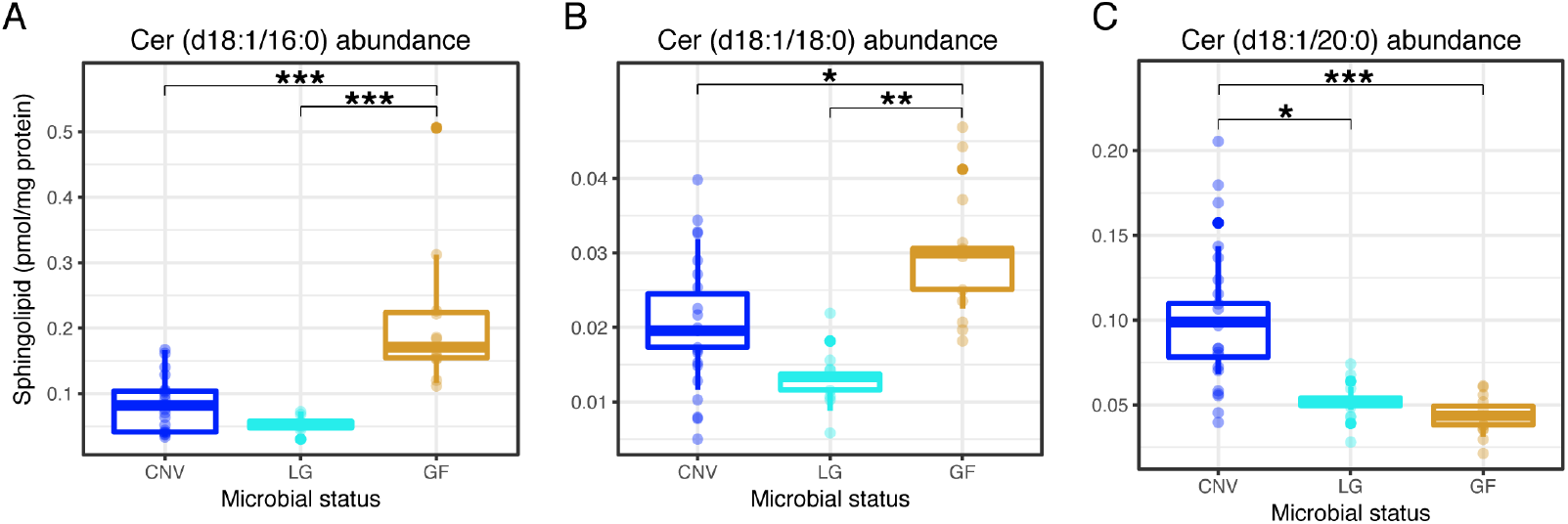
Hepatic ceramide abundances differing by microbial load Abundance of hepatic. A, Cer (d18:1/16:0), B, Cer (d18:1/18:0), and C, Cer (d18:1/20:0). Data are plotted as in Figure 5. N=8 to 21 per each microbial status group. Blue = CNV; Light blue = LG; Tan = GF.

### 4. Discussion

In the present study, we isolated the effects of the dominant oil in the Western diet consumed in the USA, where SBO makes up roughly 7% of calories. Notably, we used SBO with >50% LA, exclusively available up to about 2010, in custom HF and LF diets with SBO as the sole source of fat. This design allowed us to address whether germfree animals are resistant to increased weight gain on a diet with a greater amount of soybean oil and calories. Our results show that the fat content of the diet and associated increased caloric intake is the main driver of fat gain in the mice, regardless of the intestinal microbial load. We further show that specific circulating FAs are dependent on the gut microbiota in mice on an SBO diet. We also observe that the microbial load in the gut of SBO-fed mice impacts levels of hepatic ceramides, sphingomyelins, and sphinganine, suggesting that the gut microbiota either supply additional sphingolipids or promote hepatic ceramide and sphingomyelin synthesis.

The microbiome has been implicated in fat storage in mice. In a landmark paper, Backhed and colleagues showed that germ-free mice fed a high fat diet (with lard as a fat source) were protected from diet-induced obesity [5]. Since this early seminal observation, a more nuanced picture has emerged showing that the type of fat in the diet is an important factor in how the animal responds to a particular diet with and without a microbiome present. Germfree mice have been shown to acquire more fat mass on a high-fat compared to a low-fat diet [7–9], for instance. Two of these studies [7,9] matched the fat source between the high and low fat diet, but used a mixture of several dietary fats, including saturated fat. Our results indicate that germfree animals are not resistant to diet-induced weight gain on an SBO diet. We do observe that germfree mice acquire less adiposity and weight compared to conventional animals when matching the diets. This observation agrees with previous studies indicating that the microbiome enhances energy harvest and storage from the diet[4,5].

This study was carried out for 10 weeks to ensure diet-induced weight gain could be observed, which is challenging to do with germfree animals, especially when the diet cannot be autoclaved. One of the germfree experimental groups (GF2) was indeed contaminated at a very low level, and rather than dispose of these animals, we included them in the study as “low-germ”. The retention of this group led to the observation that liver sphingolipid levels were linearly related to microbial biomass in the gut. The addition of the LA/ALA gavage treatment at the end of the study allowed us to further ask if plasma FAs were altered specifically in the context of an acute treatment of one of the FAs in SBO, and resulted in us observing microbial interactions dependent on the gavage for C15:0 and ClnAs.

Differences in hepatic and serum lipids have been previously observed to be diet and microbiota dependent[26,29–32] and plasma lipids are well known to be directly influenced by dietary lipids[33–36]. Similar to a previous study, we observed more differential lipid molecules between conventional and germfree in the liver compared to plasma[26]. In our study, we observe many plasma FAs that differ solely by diet and/or gavage, and fewer affected by microbes. In particular, our data indicate two plasma FAs that are positively affected by the presence of microbes: C18:0 *iso* and C20:3n-6. C18:0 *iso* may be a microbial product of a branched chain amino acid and a branched short chain fatty acid[35]. C20:3n-6 (dihomo-gamma-linolenic acid, DGLA) can be made from LA by the host[36] and is a precursor of the anti-inflammatory and anti-proliferative prostaglandin PGE_1_[37–39]. Therefore, microbial modulation of DGLA may represent a mechanism by which the microbiome impacts inflammatory conditions and cancer development.

Four FAs were observed to be lower in colonized compared to germfree animals. Specially, we observed lower C15:0 in colonized mice on a HF diet, gavaged with LA compared to similar germfree mice. C15:0 is generally thought to be produced by rumen microbes, found in dairy, could be carried over in the ethanol washed casein present in the mouse diet, and it has been observed in germfree rats, dependent on diet[40]. Our observation that C15:0 is lower in HF-fed and LA-gavaged colonized animals suggests two hypotheses: (i) that gut microbes chronically conditioned to a HF-diet environment with an acute LA exposure have the capacity to metabolize C15:0; or (ii) that the absorption of C15:0 is altered, perhaps due to differences in bile acid pools[41,42] between GF and colonized mice when gavaged with LA. C14:0 was also lower in HF-CNV but not dependent on LA gavage. One ClnA (ClnA.7) was higher in GF animals only on the HF diet and another (ClnAs4) was higher in GF animals when gavaged with LA. Conjugated ALAs (ClnAs) are well known to be produced by microbes from ALA[45,46], so we are uncertain as to why our data show ClnAs increasing following LA gavage and why their levels are higher in GF animals. We also did not observe an increase in the amount of C18:0 or C18:1 with ALA gavage, which are final metabolic products in CLA and ClnA metabolism[46].

It should be noted that we previously characterized the gut microbiota composition of CNV mice fed the HF or LF SBO diets, and observed minimal to no significant differences between the microbiota composition of HF and LF diet mice[47]. Hence, we hypothesize that any diet-dependent microbiome effects on lipid pools reflect functional differences in the microbiomes of mice on the HF and LF SBO diets.

While we observed diet and microbial-dependency on plasma FAs, we observed microbial-dependency of hepatic sphingolipids. Surprisingly, hepatic sphingolipid levels were linearly related to microbial load in the cecum. In accord, the sphingolipid production capacity of the gut microbiome has recently been linked to sphingolipid levels in the liver [17]. Whether the gut microbiota act as a source of sphingolipids to the host or signal to alter host sphingolipid production and processing of sphingolipids remains to be elucidated.

The sphingoid backbone of sphingolipids is predominantly produced from palmitic acid (C16:0), which comprises 14% of the FAs in SBO and is the primary product of endogenous de novo fatty acid synthesis. Perhaps because palmitic acid is a ubiquitous component of lipids, and endogenous synthesis is downregulated on HF diets, we did not observe that animals consuming more SBO had more hepatic sphingolipids. Instead, the presence of microbes increased the amounts of specific sphingolipids. Cer (d18:1/16:0) may be an exception to this statement: the increased levels of DH Cer (d18:0/16:0) (LF diet only) and increased abundance of Cer (d18:1/16:0) may indicate increased Cer (d18:1/16:0) pools in germfree mice. Caesar et al. also noted more hepatic Cer (d18:1/16:0) in lard-fed germfree compared to conventional mice[26]. Our data using SBO fed mice, differ from their observations of lower levels of Cer (d18:1/20:0) and Cer (d18:1/24:1) in conventional animals on a lard or fish oil diet[26]. Rather we observed these ceramides were lowest in our germfree mice. Finally, increased levels of hepatic Sa (d18:0) have been associated with increased mitochondrial respiration and obesity[48]. As we observed greater hepatic Sa (d18:0) in conventional mice, these data may signal increased hepatic mitochondrial function and be linked to the greater absolute weight of conventional animals.

Our findings indicate that a diet high in SBO leads to fat gain in mice regardless of the presence of a microbiome, although colonized mice were able to accumulate more fat. The microbiome density in the gut has a greater effect that dietary fat content on the sphingolipids in the liver. Collectively, these results suggest that the microbiome is involved in the alteration of host signaling related to both lipid processing and lipid storage.

## Author contributions

**Sara C. Di Rienzi:** Conceptualization, Data curation, Formal analysis, Funding acquisition, Investigation, Methodology, Software, Visualization, Writing – original draft, Writing - review & editing; **Elizabeth L. Johnson:** Conceptualization, Data curation, Formal analysis, Investigation, Methodology, Writing - review & editing; **Elizabeth A. Kennedy**: Data curation, Formal analysis, Investigation; **Juliet Jacobson**: Data curation, Formal analysis, Investigation; **Peter Lawrence**: Data curation, Formal analysis; **Dong Hao Wang**: Data curation, Formal analysis; **Tilla S. Worgall**: Resources, Supervision; **J. Thomas Brenna**: Conceptualization, Resources, Supervision; Writing - review & editing; **Ruth E. Ley**: Resources, Writing - review & editing, Supervision, Project administration, Funding acquisition.

## Abbreviations

(ALA): alpha-linolenic acid
(ceramide): Cer
(CLA): conjugated linoleic acid
(ClnA): conjugated linolenic acid
(CNV): conventional
(DGLA): dihomo-gamma-linolenic acid
(DH): dihydro
(EPA): eicosapentaenoic acid
(FA): fatty acid
(GF): germfree
(HF): high fat
(LA): linoleic acid
(LF): low fat
(LG): low-germ
(MS): microbial status
(PCoA): Principal Coordinate Analysis
(Sa): sphinganine
(SBO): soybean oil
(SM): sphingomyelin
(So): sphingosine

## Acknowledgements/grant support

We thank members of the Ley lab, Jiyao Zhang, the staff of the Cornell Animal Facility, Jennifer Mosher, Sylvie Allen, and Lynn Marie Johnson for their assistance, helpful discussions, and insight. S.C.D.R. is an Eli & Edythe Broad Fellow of the Life Sciences Research Foundation. This work was funded by NIH grant DP2OD007444 and by the Max Planck Society. The content is solely the responsibility of the authors and does not necessarily represent the official views of the National Institutes of Health.

## Supplementary Material

### Supplementary Tables 1-3

**Supplementary Table 1**. Least squares means estimates of significant experimental variables on plasma fatty acids.

**Supplementary Table 2**. Least squares means estimates of significant experimental variables on hepatic sphingolipids.

**Supplementary Table 3**. Least squares means estimates of significant experimental variables on hepatic ceramide abundance.

### Supplementary Figures S1-6: Legends

**Figure S1:** Plasma FAs C14:0 and C16:0 *iso* differ by diet, gavage, and/or microbial status. Data are plotted as in Figure 5. N=3 to 148 per group.

**Figure S2:** Plasma FAs C15:0, C16:0, and C16:1n-7 differ by diet, gavage, and/or microbial status. Data are plotted as in Figure 5. N=3 to 148 per group.

**Figure S3:** Plasma FAs C18:0 *iso*, C18:1n-7/9/?, and C18:2n-6 differ by diet, gavage, and/or microbial status. Data are plotted as in Figure 5. N=3 to 148 per group.

**Figure S4:** Plasma FAs C18:3n-6, C18:3n-3, and C20:3n-6 differ by diet, gavage, and/or microbial status. Data are plotted as in Figure 5. N=3 to 148 per group.

**Figure S5:** Plasma FAs ClnA.7 and C22:5n-6 differ by diet, gavage, and/or microbial status. Data are plotted as in Figure 5. N=3 to 148 per group.

**Figure S6:** Plasma FAs C20:5n-3, C24:1n-9, ClnAs4, and ClnA.2 differ by diet, gavage, and/or microbial status. Data are plotted as in Figure 5. N=3 to 148 per group.

