## Supplementary figures and images for "The microbiome affects liver sphingolipids and plasma fatty acids in a murine model of the Western diet based on soybean oil"

### Supplemental figures

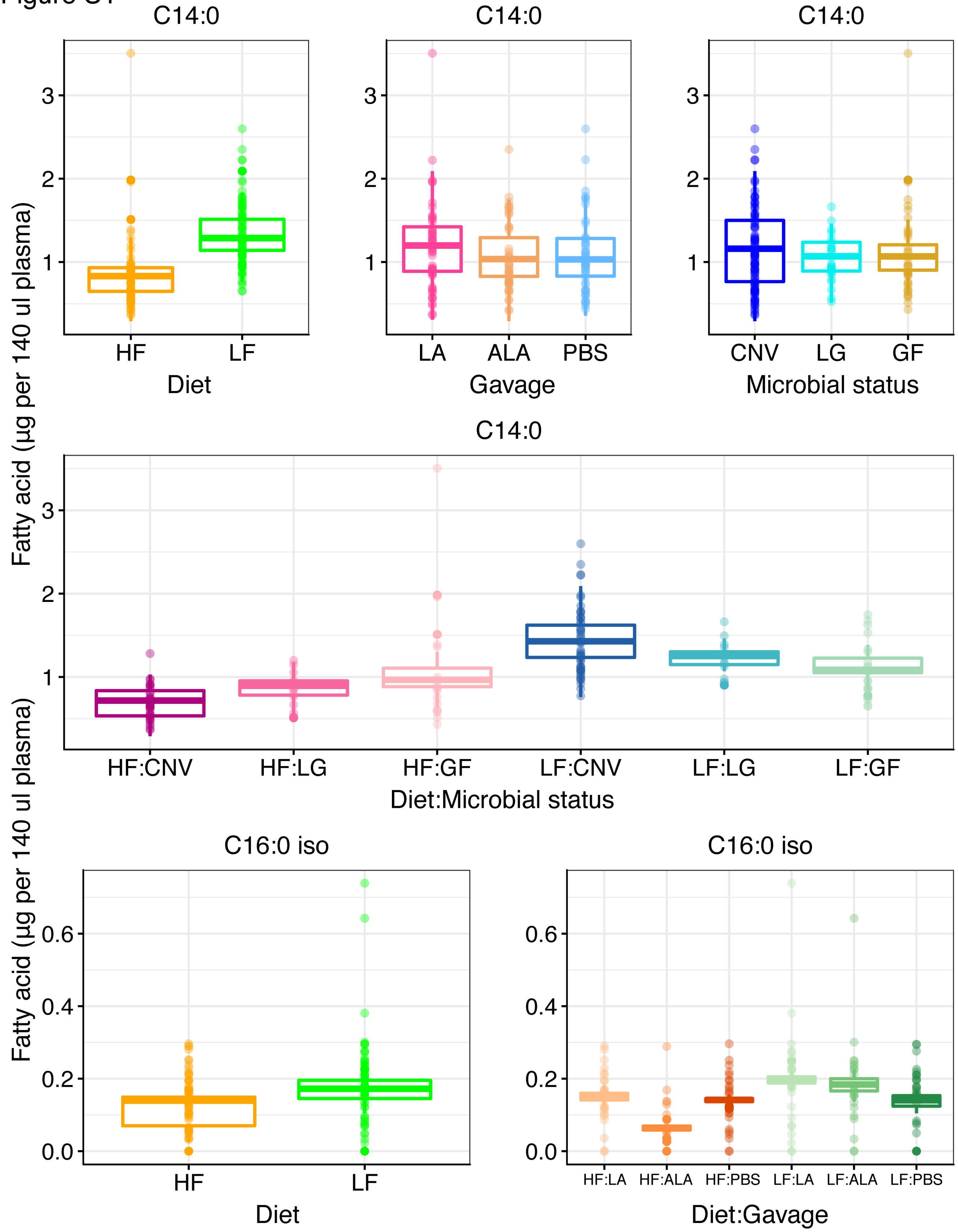

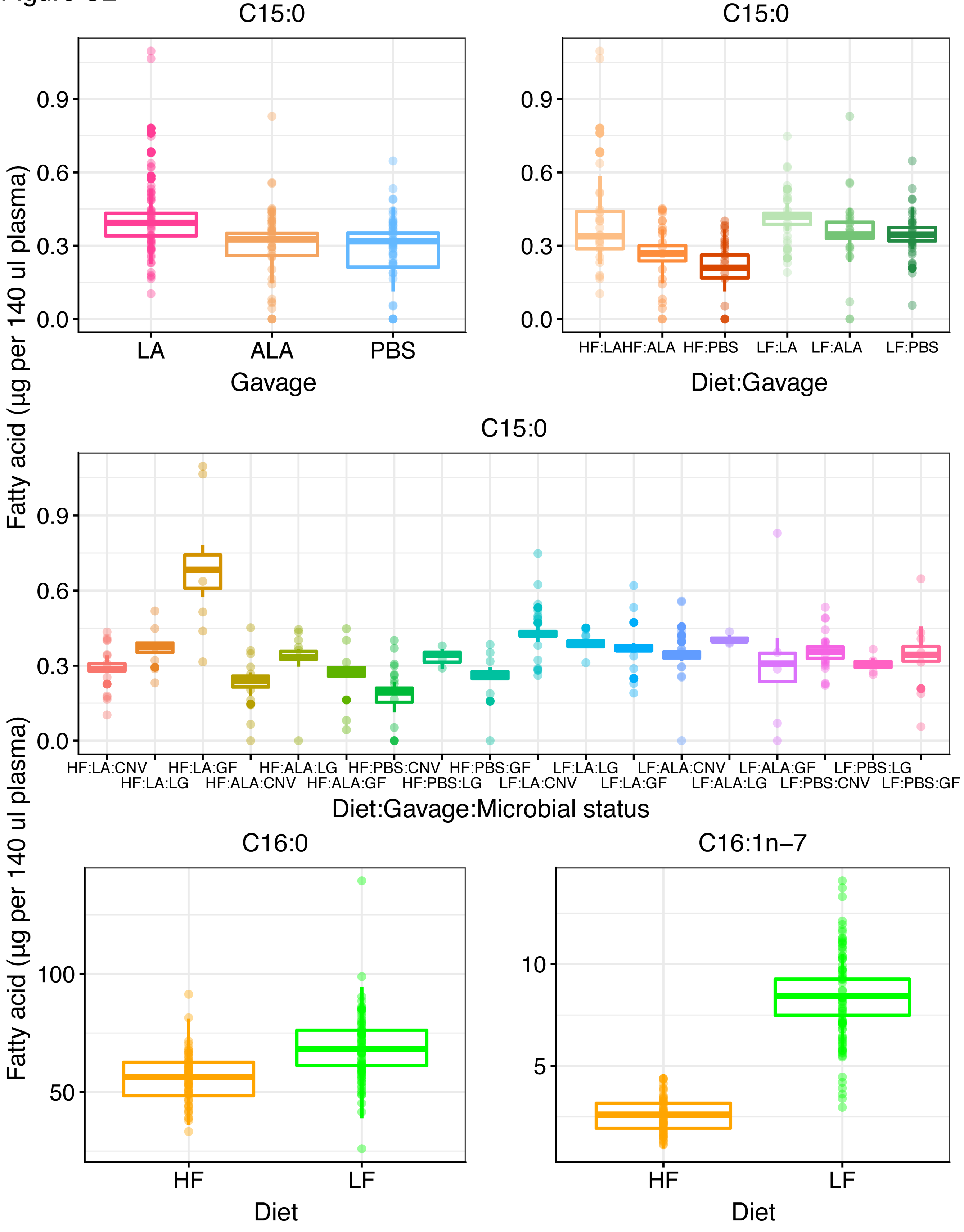

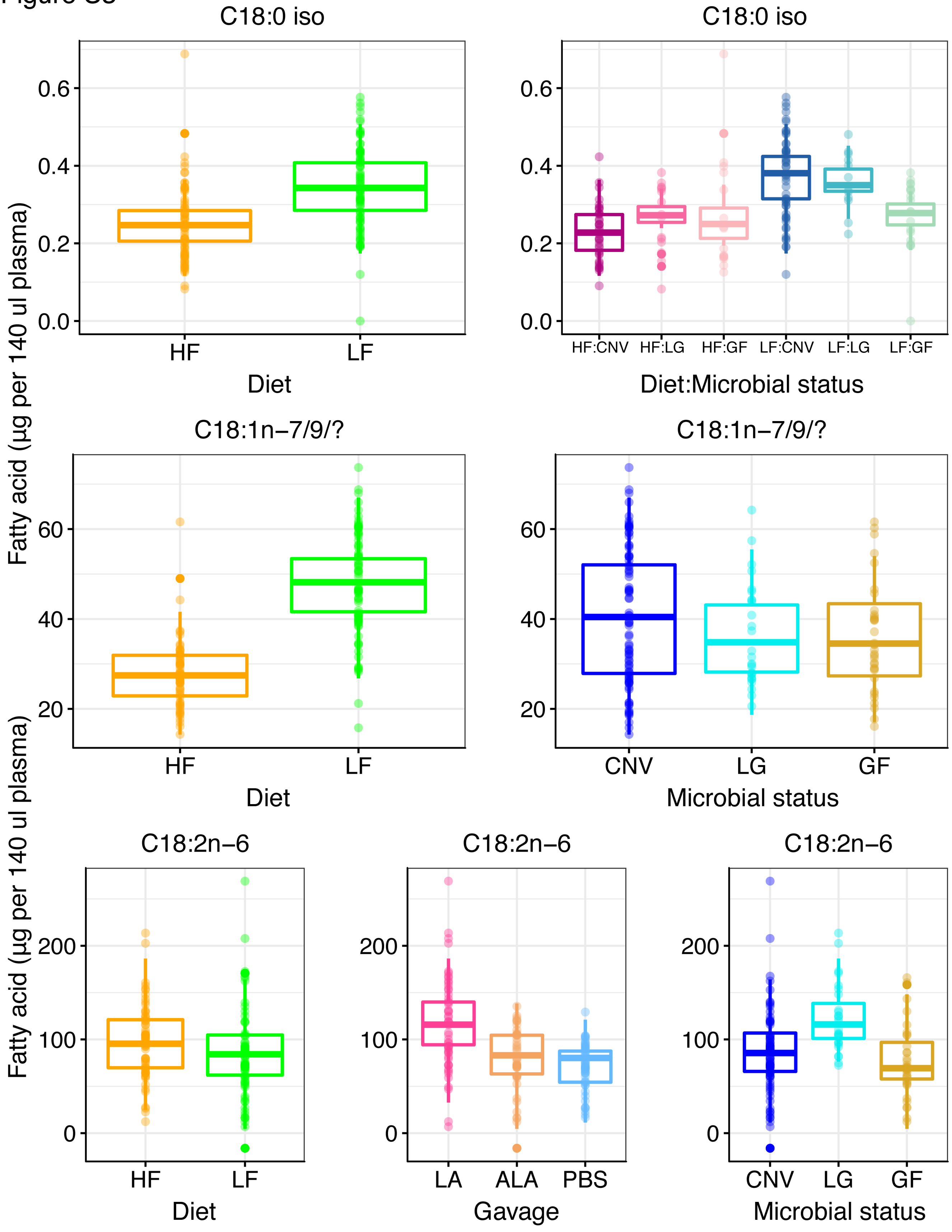

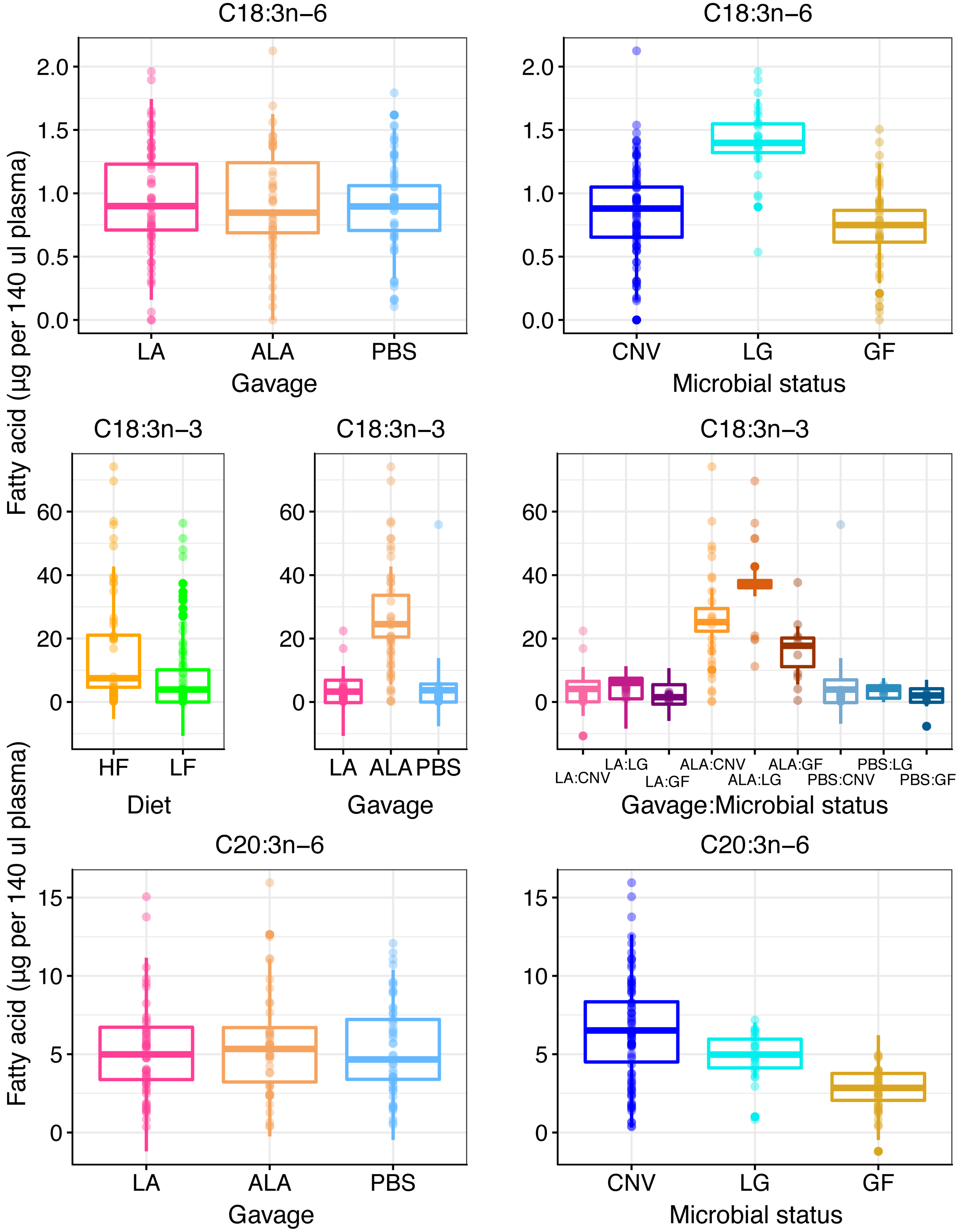

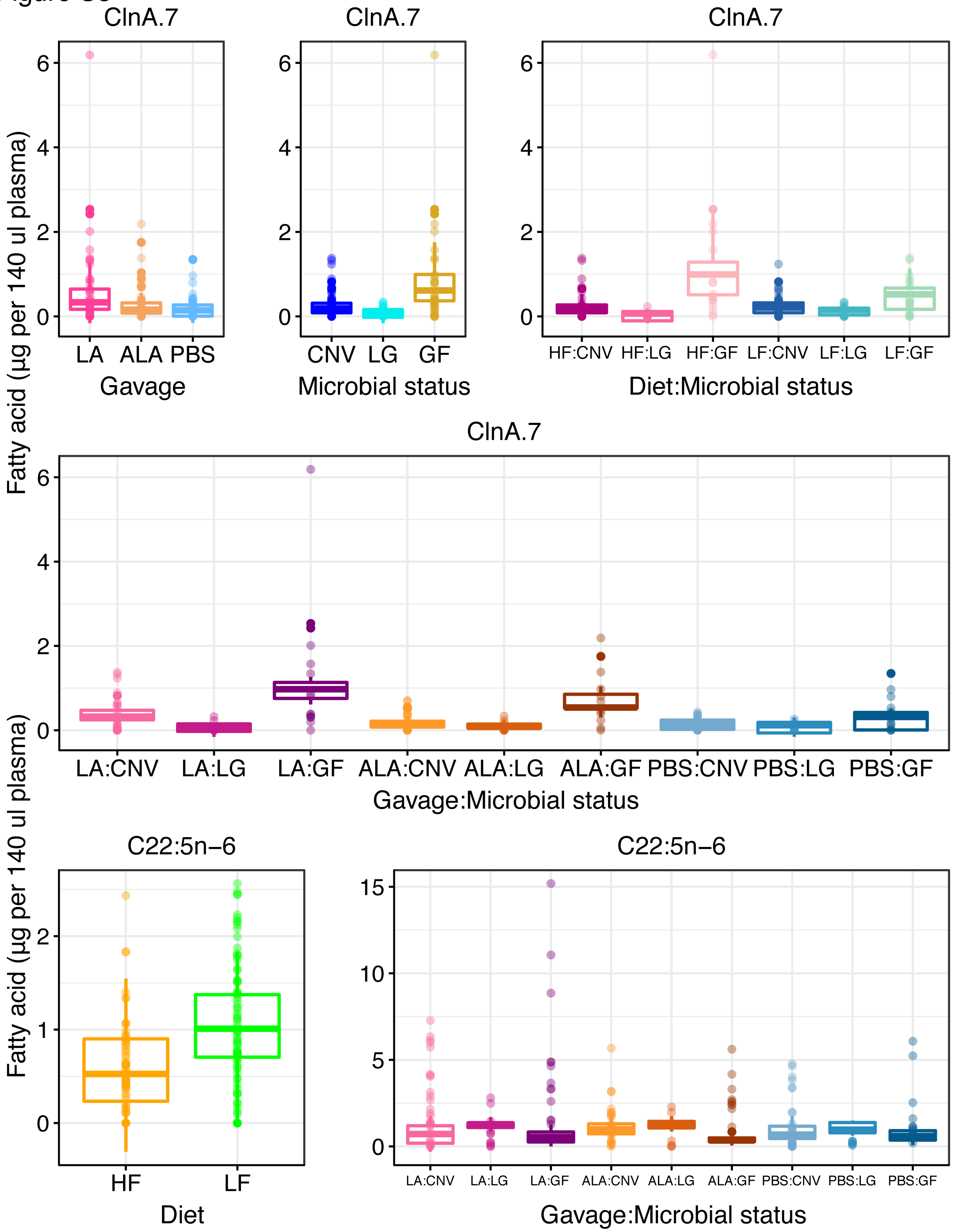

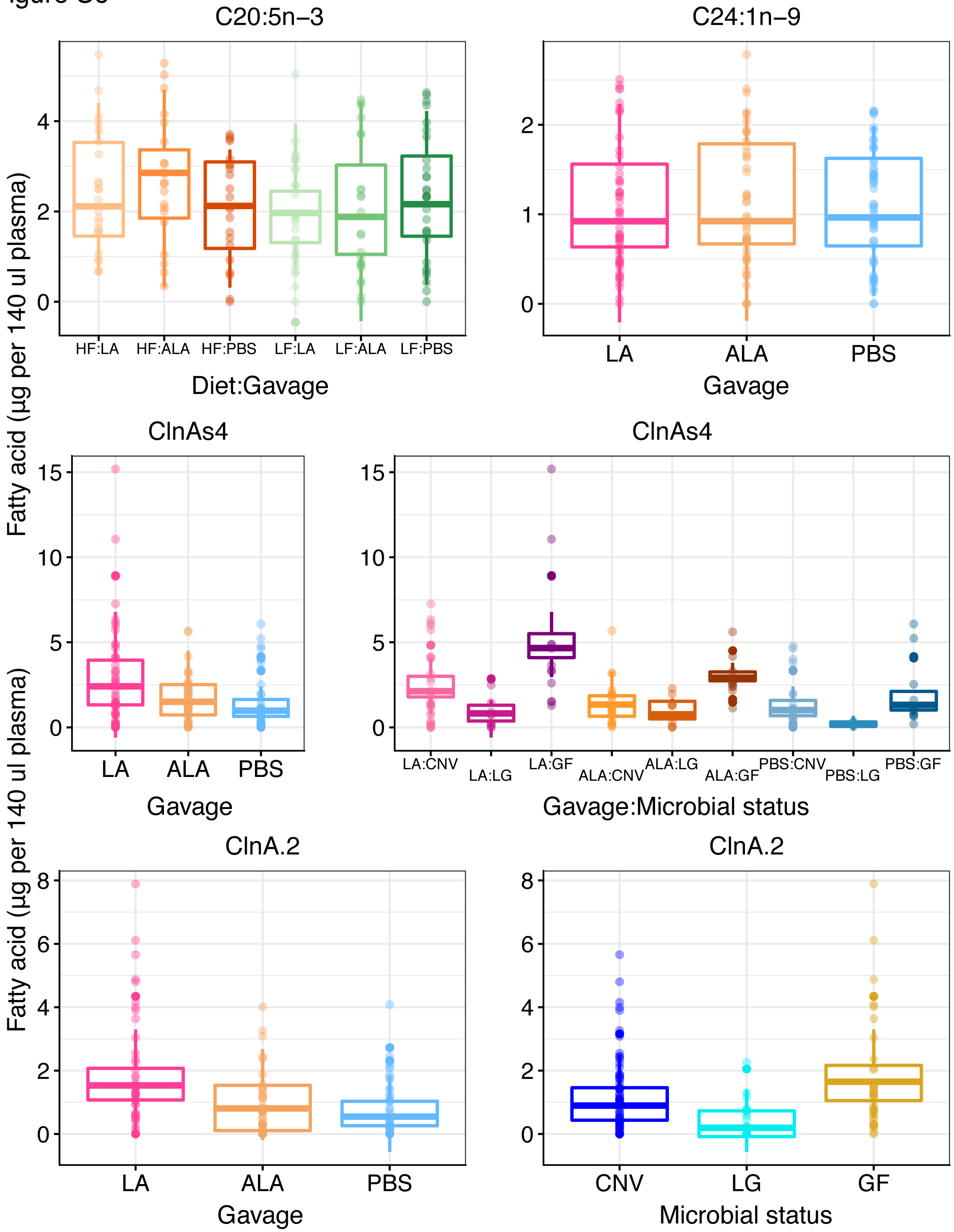
